# Effect of Increasing Glucose Concentration on Estimation of Electrolytes by ion selective electrode

**DOI:** 10.1101/2020.11.25.396440

**Authors:** Omalkhair A. Khidir, Mohammed Elsheikh Osman

## Abstract

**Background:** The measurement of serum electrolytes like sodium (Na+), potassium (K+) is routinely performed in clinical biochemistry laboratories using ion-selective electrodes (ISE).

**Aim:** To minimize error and test the effect of increasing glucose concentration on estimation of sodium and potassium.

**Materials and Methods:** Randomly selected sera samples submitted for routine biochemical estimations were pooled for the study. A stock solution of glucose with concentration of 20 g/ dL was prepared by dissolving anhydrous glucose in normal saline (0.9 % NaCl). Next the pooled sera was divided into several aliquots with 1.0 ml sera each and increasing amounts of the stock solution were added to the different aliquots so as to generate glucose concentrations ranging from around 100 mg/dL to about 3000 mg/dL.

**Results:** the glucose concentration of different serum aliquots obtained on cobas c311(roche system) were found to rang from 110 to 2577 mg/dl. The mean of sodium by direct ion selective 142±0.52 mmol/l, values showed in significant difference (p=0.320).The mean of sodium by indirect ion selective 143 ± 0.72 mmol/l, values showed significant difference (p< 0.05). And the mean of potassium by direct ion selective 3.54 ±0.29 mmol/l, values showed significant difference (p< 0.05).

The mean of potassium by indirect ion selective 3.53±0.26 mmol/l, values showed significant difference (p< 0.05).

**Conclusions:** Increasing blood gluose interferance in electrlytes mainly potasssium by both direct and indirect ion selective electrode, whenincreasing glucose potassium decreae significantly No different significant between direct and indirect ion selective electrods in potassium when used cobas c311 and easy lyte.Significant interferance in sodium by indirect ion selective electrodewhile no significant interferance in sodium direct ion selective electrode.Glucose concentration interfer the electrolyte specially potassium in concentration more than 1700 mg/dl by direct and indirect methods.

## Introduction

The measurement of serum electrolytes like sodium (Na+), potassium (K+) is routinely performed in clinical biochemistry laboratories using ion-selective electrodes (ISE). An ISE generates a difference in electrical potential between itself and a reference electrode when the cell current is zero i.e. at equilibrium. The ISE membrane is the key component of all potentiometric ion sensors and it is composition determines the optimal selectivity to the ion of interest. Specific ISE membranes can be made of glass, crystals or some specific ionophore may be incorporated in the matrix, which determines selectivity of the electrode. The membrane potential caused by selective permeability of a membrane to a particular ion and the potential generated at membrane—test solution interface is proportional to the log of ionic activity or concentration of the selected ion in solution as expressed by the Nernst equation [1].

Two methods have been described for measurement of serum electrolytes by ISE: direct and indirect. The initial methods (Indirect ISE) were based on dilution of serum with a buffer, as in flame photometry, and specimen is brought to the electrode surface without dilution and activity of the relevant ion is measured in the plasma water. The dilution of a sample has important implications in cases where the solid component of plasma is increased and is known as ‘electrolyte exclusion effect’ which often leads to falsely low values of serum electrolytes especially Na+, recognized as ‘pseudohyponatremia’[3].

This arises most commonly in situations of extreme hyperproteinemia or hyperlipidemia. Numerous methods have been employed by various authors to measure corrected Na+, K+ and Cl-based on the concentration of protein, albumin or triglycerides. However, they are prone to either over – estimation of fall in sodium levels or are complex calculations or do not take into account all the parameters comprising solid phase of plasma [4]. Direct ISE methods are not affected by this phenomenon as there is no dilution of the sample and moreover, whole blood can be used directly for rapid estimation as in case of open heart surgery. The electrolyte exclu – sion effect not only affects sodium but also other ions like potassium, chloride, magnesium, bicarbonate etc. However reduction in sodium often becomes clinically significant as it falls below the reference range due to this effect. Hence, the term pseudohyponatremia has gained more acceptance. Besides protein and triglycerides, other solutes like glucose, urea etc. might also affect the analysis of serum electrolytes by direct and indirect potentiometric methods. One such case has been reported in literature where in the authors have shown discrepancies between the two methods with high glucose concentrations which resolved with subsequent treatment and fall in glucose levels [5].

The present study was conducted with an aim to study the effect of increasing glucose concentrations on estimation of electrolytes like Na+, K+by direct as well as indirect ISE methods.

### Analytical Methodology for the Determination of sodium and 1-1 Potassium

Sodium is determined by atomic absorption spectrophotometry (AAS), flame emission spectrophotometry(FES), electrochemically with a Na+-ISE, or spectrophotometric methods all have been used for Na+ and K+ analysis. Most laboratories, however, now use ISE methods. For example, of the more than 5000 laboratories reporting College of American Pathologists (CAP) proficiency survey data for Na+ and K+, >99% were using ISE methods in2005. The principles of each of these approaches are the same whether the instrumentation is dedicated or integrated into a multichannel system. The electrolyte exclusion effect also will affect the measurement of Na+ and K+ [1].

### Ion-Selective Electrodes: 1-2

An ISE is a special-purpose, potentiometric electrode consisting of species. The potential pduced at the membrane-sample solution interface is proportional to the logarithm of the ionic activity or concentration.ISEs integrated into chemical analyzers usually containNa+ electrodes with glass membranes and K+ electrodes with liquid ion-exchange membranes that incorporate valinomycin. In practice, apotentiom-etric measuring system is calibrated by introduction of calibrator solutions containing defined amounts of Na+ and K+. The potentials of the calibrators are determined, and the AE/A log concentration is stored in computer memory as a factor for calculating unknown concentration when E of the unknown is measured.

Frequent calibration, initiated either by the user or by automated uptake of sample from a reservoir of calibrator,is characteristic of most systems. Some instruments are designed to measure Na+ and K+ in whole blood, particularly point of care testing devices and newer blood gas analyzers. Indirect ISE, sample is introduced into the measurement chamber Indirect and direct are the two types of ISEs. With an after mixing with a large volume of diluent. This use of a larger volume is advantageous because it adequately covers the surface of a large electrode and minimizes the concentration of protein at the electrode surface. lndirect ISEs are most common in large, high-throughput automated analyzers. In the common in large, high-throughput automated analyzers. in the direct ISE methods, sample is presented to the electrodes without dilution Direct ISEs are used on blood gas analyzers, point of care devices, and Other single-use instruments. A 2005 CAP proficiency testing report indicated that approximately two thirds of the laboratories used indirect ISE proficiency to measure Na+ and K+. Important differences in direct And indirect methods that cause significant differences in analytical results are discussed in the later section on the electrolyte exclusion effect Errors observed in the use of ISEs are due to:

1. lack of analytical selectivity,
2. repeated protein coating of the ionsensitive membrane, and
3. contamination of the membrane

or salt bridee by, ions that compete or react with the selected ion and thus alter electrode response. These errors necessitate periodic changes of the membrane as part of routine maintenance [1].

### Electrolyte Exclusion Effect: 1-3

The electrolyte exclusion effect is the exclusion of electrolytes from the fraction of the total plasma volume that is occupied by solids. The volume of total solids (primarily protein and lipid) in an aliquot of plasma is approximately 7%. Thus −93% of plasma volume is actually water. The main electrolytes (Na+, K+, C1-, HCO3-) are essentially confined to the water phase. When a fixed volume of total plasma,for example 10 uL, is pipetted for dilution before flame photometry or indirect ISE analysis, only 9.3 uL of plasma water containing the electrolytes is actually added to the diluent. Thus a concentration of Na+ determined by flame photometry or indirect ISE to be 145 mmol/L is the concentration in the total plasma volume, not in the plasma water volume. In fact, if the plasma contains 93% water, the concentration of Na+ in plasma water is 145 x (100/93), or 156 mmolL. This negative “error” in plasma electrolyte analysis has been recognized for many years. Even though it is the electrolyte concentration in plasma water that is physiological, it was assumed that the volume fraction of water in plasma is sufficiently constant that this difference could be ignored. In fact, all electrolyte reference intervals are based on this assumption and actually reflect concentrations in total plasma volume and not in the water volume. Indeed, virtually all concentrations measured in the clinical chemistry laboratory are related to the total sample volume rather than to the water volume. This electrolyte exclusion effect becomes problematic when pathophysiological conditions are present that alter the plasma water volume, such as hyperlipidemia or hyperproteinemia. In these settings, falsely low electrolyte values are obtained whenever samples are diluted before analysis, as in flame photometry or indirect ISE. It is the dilution of total plasma volume and the assumption that plasma Water volume is constant that renders both indirect ISE and flame – photometry methods equally subject to the electrolyte exclusion effect. In certain settings, such as ketoacidosis with severe hyperlipidemia or multiple myeloma with severe hyperproteinemia, the negative exclusion effect may be so large that laboratory results lead clinicians to believe that electrolyte concentrations are normal or low when, in fact, the concentration in the water phase may be high or normal, respectively.’ In direct ISE methods, there is no dilution and measured electrolyte activity is directly proportional to the concentration in the water phase, not the concentration in the total volume. To make results from direct ISEs equivalent to flame photometry and indirect ISEs, most direct ISEs, most direct ISE methods operate in what is commonly referred to as the “flame mode.” In this mode, the directly measured concentration in plasma water is multiplied by the average water volume fraction of plasma(0.93). Although the latter may vary widely, as long as the activity of the specific ion is constant, the concentration of the ion in the water phase becomes independent of the relative proportions of water and total solids if the ion is not bound by proteins, as is the case for Ca++. Therefore direct ISE methods are free of the electrolyte exclusion effects, and the values determined by direct ISE methods, even in the flame mode, are directly proportional to activity in the water phase and define electrolyte concentrations in a more physiological and physicoch-ernical sense. Direct ISE methods are now considered as the methods of choice for electrolyte analysis. This is based on the fact that large changes in plasma lipid, protein, and other solids often occur in relatively common clinical conditions [1].

### 1-4 increasing blood glucose

According to the World Diabetes Foundation, diabetes is the worlds fastest growing chronic disease that affects 6.4% of the world’s adult population [10],The accuracy of the clinical laboratory tests is important for patient care, and control of the whole testing process is the responsibility of laboratory professionals. Though the errors in the analytical phase are under strict control by the improved technology and control materials, body fluid compounds are known to interfere with some analytical reaction steps that are not yet solved by the manufacturers [6] The major endogenous substances that interfere with the laboratory analyses are hemoglobin, bilirubin, and lipids Because the magnitude of interference varies from method to method and according to the concentration of the interference [7,9], we aimed to experiment the interfering effect of glucose at different levels on measure Na+ and K+.

### 5 Rationale

Interferences like proteins, triglycerides, drugs etc. are known to affect in estimation of sodium and potassium. The present study was designed to look into the effect of increasing glucose concentrations on estimation of Na, K.

Because the magnitude of interference varies from method to method method and according to the concentration of the interference we aimed to experiment the interfering effect of glucose at different levels on estimation electrolyte.

This study will help authorities to evaluate the problem more objectively and implement appropriate measures of electrolytes.

## 2 Objectives

### 2-1 General objective

To calculate the Effect of Increasing Glucose Concentrations on Estimation of Electrolytes by ion selective electrode.

### 2 Specific objectives

1. To calculate the effect of Increasing Glucose Concentrations on estimate of serum sodium by direct ion selective electrode.
2. To calculate the effect of Increasing Glucose Concentrations on estimate of serum sodium by indirect ion selective electrode.
3. To calculate the effect of Increasing Glucose Concentrations on estimate of serum potassium by direct ion selective electrode.
4. To calculate the effect of Increasing Glucose Concentrations on estimate serum potassium by indirect ion selective electrode.
5. Correlation between direct and indirect ion selective electrode.

### 3 Material and methods

#### 3-1 Preparation of Samples

(interference study)

Randomly selected sera samples submitted for routine biochemical estimations were pooled for the study. The baseline data for the electrolytes, glucose, liver function tests, renal function tests, total protein, albumin and triglycerides were run on Cobas c311 analyzer Daily Prep and maintenance – Roche Diagnostics USA. after routine calibration and quality control check. Mean for A stock solution of glucose with concentration of 20 g/ dL was prepared by dissolving anhydrous glucose in normal saline (0.9 % NaCl). Next the pooled sera was divided into several aliquots with 1.0 ml sera each and increasing amounts of the stock solution were added to the different aliquots so as to generate glucose concentrations ranging from around 100 mg/dL to about 3000 mg/dL; volumes of the different aliquots were made up to 1.0 ml.

#### 3.2 Inclusion criteria

There was no involvement of any direct human or animal subjects in any experiments in the study. All procedures performed in the study involving samples from human origin.

#### 3.3 Exclusion criteria

1. Samples from Patients who received chemotherapy, radiotherapy or hormonal therapy.
2. Samples from Patients with history or hyper protenimia and hyper triglyceridimia
3. Hemolysis and bilirubin in sample
4. Samples from diabetic patients.

#### 3-4 Measurement of analyses

Serum was test by both direct and indirect ion selective electrode Analyzer, Easy lyte full automated for Direct ISE and Cobas c311 auto Analyzer for Indirect ISE.

Glucose concentration was measured by Cobas c311 auto analyzer using glucose oxidase method.

#### 3-5 Ethical Standards

All procedures performed in the study involving samples from human origin were in accordance with the ethical standards of the institutional and / or national research committee.

#### 3-6 Quality Control

The precision and accuracy of all methods used in the study has been checked each time a batch was analyzed by including commercially prepared control sera.

#### 3-7 statistic anaylisis

Results are presented as mean ± SD values and two-way ANOVA with Bonferroni post-test was performed using Prism5TM software to analyse the differences between direct and indirect ISE values and also at different glucose concentrations with respect to the baseline value. Pearson’s correlation was calculated using SPSS Version 21. P<0.05 was considered to be statistically significant.

## 4 Results

### Baseline biochemistry profile: 4-1

The total protein and triglyceride concentration were observed to be within physiological range in the pooled sera ruling out any significant interference due to these anaytes.

The glucose concentration of different serum aliquots obtained on cobas c311 (roche system) were estimated after addition of glucose solution and found the range from 110 to 2577 mg/dl.

### Analysis of electrolytes by direct and indirect ISE Methods: 4-2

The mean of sodium by direct ion selective 142±0.52 mmol/l, values showed in significant difference (p=0.320).

The mean of sodium by indirect ion selective 143 ± 0.72 mmol/l, values showed significant difference (p< 0.05).

And the mean of potassium by direct ion selective 3.54 ±0.29 mmol/l, values showed significant difference (p< 0.05).

The mean of potassium by indirect ion selective 3.53±0.26 mmol/l, values showed significant difference (p< 0.05).

The level of Na + and K+ in serum at different glucose concentration as measured by direct and indirect ISE. K+ values by direct and indirect showed significant difference (p<0.05) with difference glucose concentration Fig 2,4.

However, no significant difference was observed in Na+ by Direct Methods Fig 1.

Significant difference was observed in Na+ by in Direct methods Fig 3.

### Correlation Between Direct and indirect ISE: 4-3

The persons correlation coefficient, R was was estimated for each of the electrolytes for both direct and indirect ISE with glucose concentration the K+values showed strong negative correlation by direct and indirect R= (−0.998). Significant correlation was showed in Na+ by indirect ISE (R=0.730).

**Figure No(4-1).**
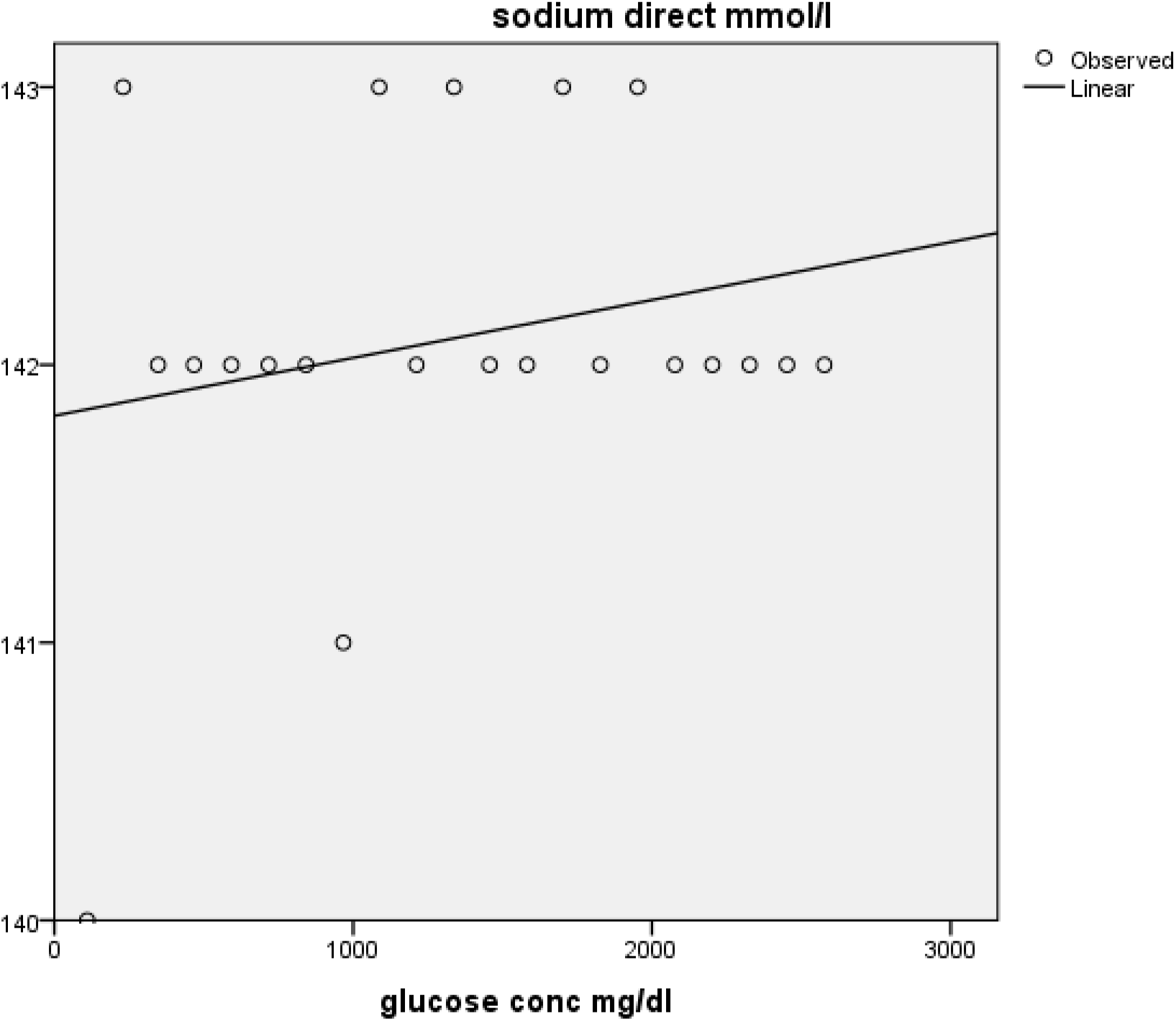
The effect of Increasing glucose concentration in sodium by direct ion selective electrode. R square = 0.052 Pvalue = 0.320

**Figure No(4-2).**
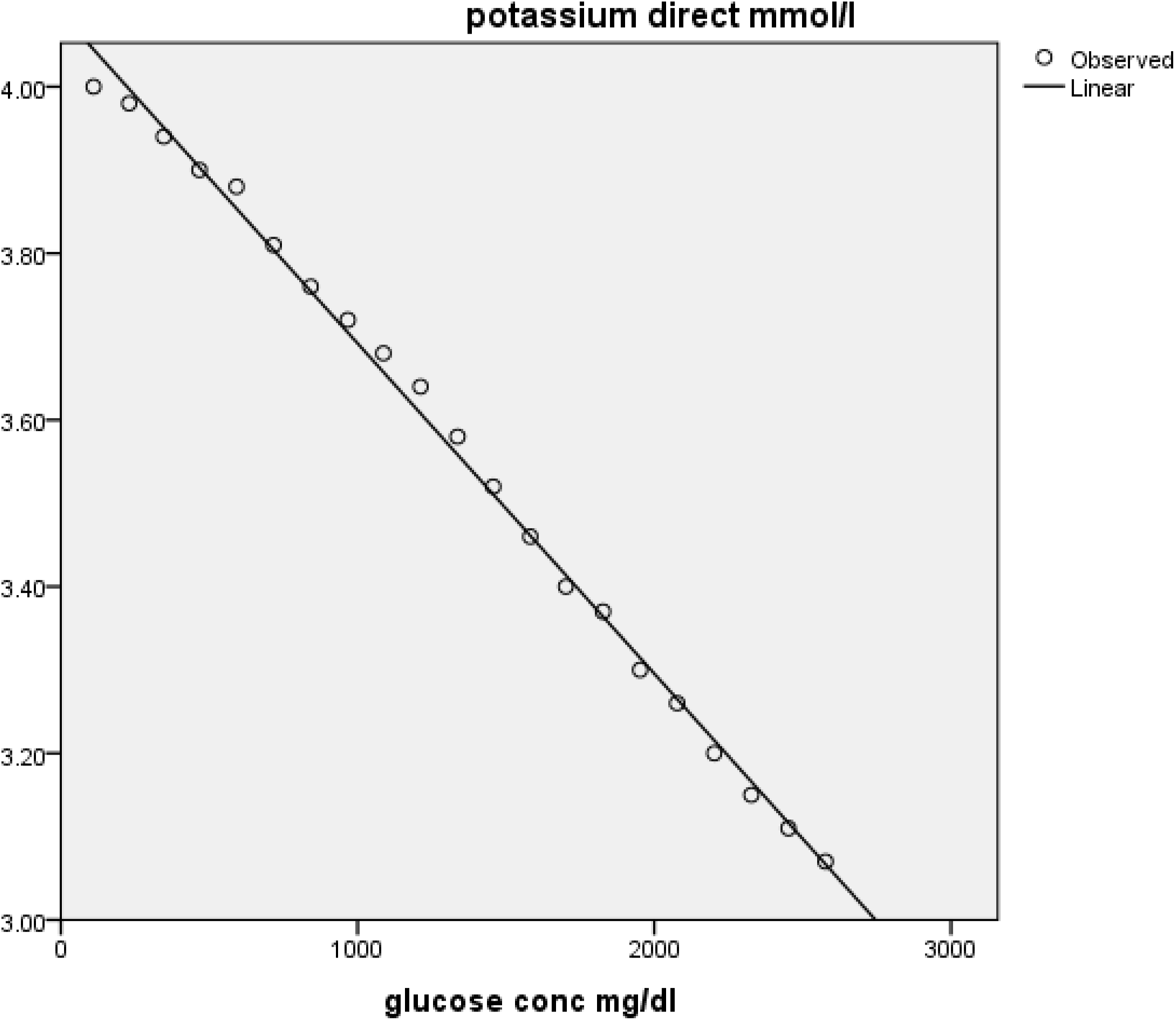
The effect of Increasing glucose concentration in potassium by direct ion selective electrode. R square = 0.996 Pvalue = 0.000

**Figure No(4-3).**
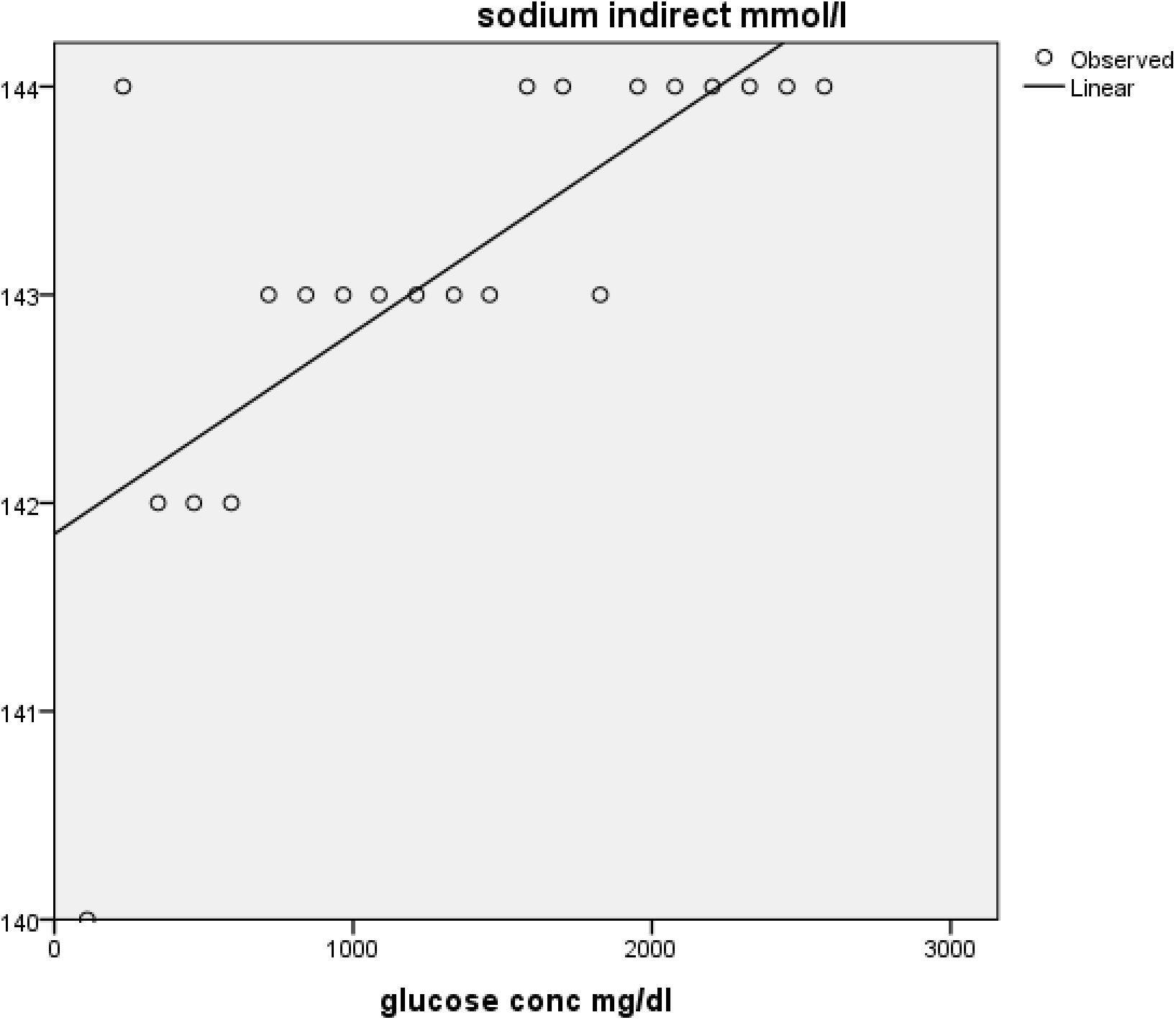
The effect of Increasing glucose concentration in sodium by indirect ion selective electrode. R square = 0.534 Pvalue = 0.000

**Figure No(4-4).**
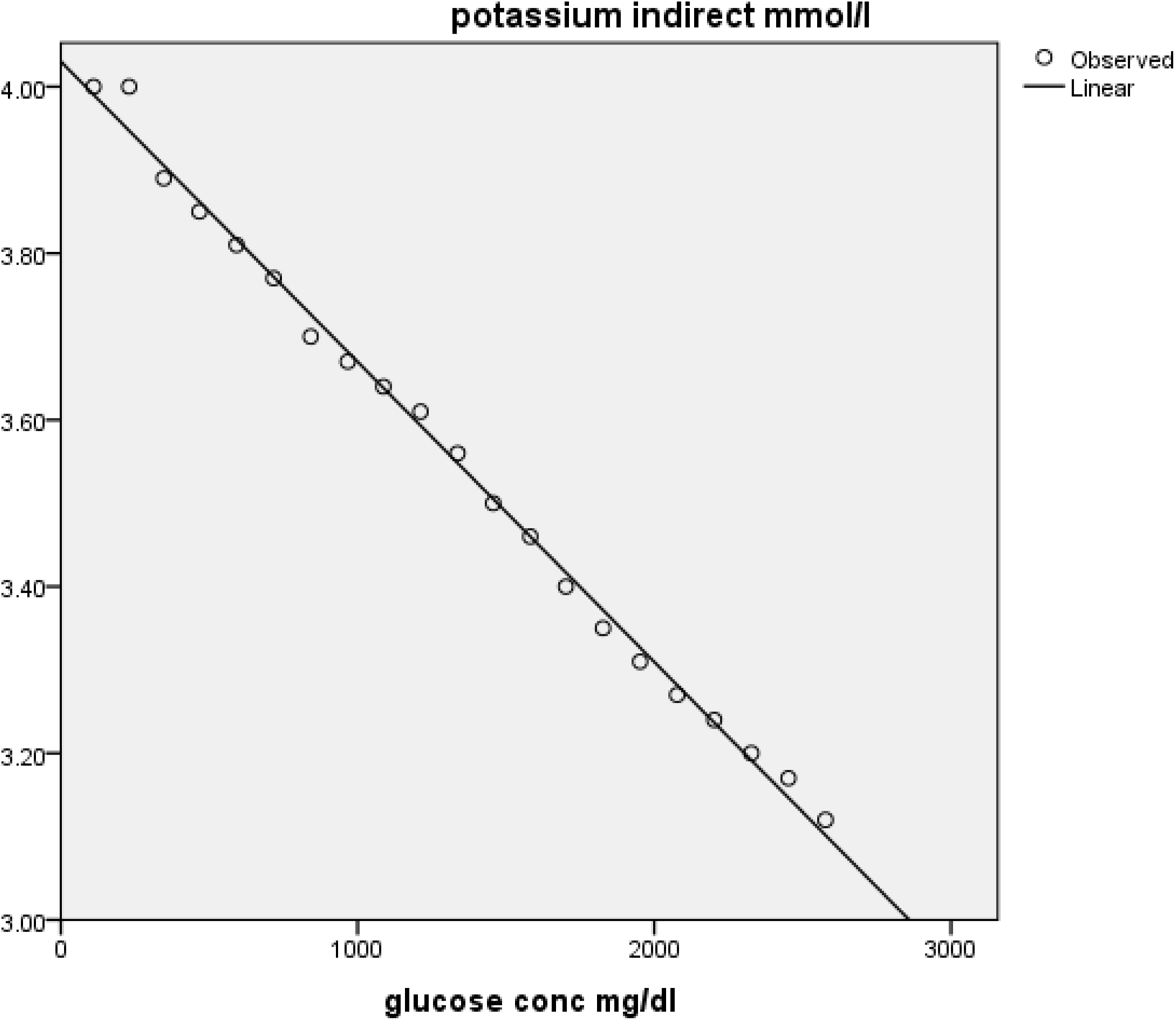
The effect of Increasing glucose concentration in potassium by indirect ion selective electrode. R square = 0.996 Pvalue = 0.000

### 4-4 Study for interferance

Table NO (4-4-1): Group 1 glucose concentration less than 1000 mg/dl and error in sodium, potassium.

(Allowable error Na+ 4mmol/l, K+ 0.5 mmol/l)

| No. | Profile | Glucose conc. | Na<sup>+</sup> direct | K<sup>+</sup> direct | Na<sup>+</sup> direct | K<sup>+</sup> indirect |
| --- | --- | --- | --- | --- | --- | --- |
| 1 | Test<br>Baseline<br>Error<br>Allowable Error | 230 | 143<br>140<br>3<br>acceptable | 3.98<br>3.93<br>0.05<br>acceptable | 144<br>140<br>4<br>acceptable | 4.0<br>3.93<br>0.07<br>acceptable |
| 2 | Test<br>Baseline<br>Error<br>Allowable Error | 347 | 142<br>140<br>2<br>acceptable | 3.94<br>3.93<br>0.01<br>acceptable | 142<br>140<br>2<br>acceptable | 3.89<br>3.93<br>-0.04<br>acceptable |
| 3 | Test<br>Baseline | 467 | 142<br>140 | 3.90<br>3.93 | 142<br>140 | 3.85<br>3.93 |
|  | Error Allowable Error |  | 2<br>acceptable | -0.03<br>acceptable | 2<br>acceptable | -.008<br>acceptable |
| 4 | Test<br>Baseline<br>Error<br>Allowable Error | 592 | 142<br>140<br>2<br>acceptable | 3.88<br>3.93<br>-0.05<br>acceptable | 142<br>140<br>2<br>acceptable | 3.81<br>3.93<br>-0.12<br>acceptable |
| 5 | Test<br>Baseline<br>Error<br>Allowable Error | 717 | 142<br>140<br>2<br>acceptable | 3.81<br>3.93<br>-0.12<br>acceptable | 143<br>140<br>3<br>acceptable | 3.77<br>3.93<br>-0.16<br>acceptable |
| 6 | Test<br>Baseline<br>Error<br>Allowable Error | 842 | 142<br>140<br>2<br>acceptable | 3.76<br>3.93<br>- 0.17<br>acceptable | 143<br>140<br>3<br>acceptable | 3.70<br>3.93<br>-0.23<br>acceptable |
| 7 | Test<br>Baseline<br>Error<br>Allowable Error | 967 | 141<br>140<br>1<br>acceptable | 3.72<br>3.93<br>-0.21<br>acceptable | 143<br>140<br>3<br>acceptable | 3.67<br>3.93<br>-0.26<br>acceptable |

Table NO (4-4-2): Group 2 glucose concentration from 1000 to 2000 mg/dl and error in sodium, potassium.

(Allowable error Na+ 4mmol/l, K+ 0.5 mmol/l)

Table NO (4-4-2) :Group 2 glucose concentration from 1000 to 2000 mg/dl and error in sodium ,potassium. (Allowable error Na+ 4mmol/l ,K+ 0.5 mmol/l )
| No. | Profile | Glucose conc. | Na+ direct | K+ direct | Na+ direct | K+ indirect |
| --- | --- | --- | --- | --- | --- | --- |
| 1 | Test<br>Baseline<br>Error<br>Allowable Error | 1087 | 143<br>140<br>3<br>acceptable | 3.68<br>3.93<br>0.25<br>acceptable | 143<br>140<br>3<br>acceptable | 3.64<br>3.93<br>-0.29<br>acceptable |
| 2 | Test<br>Baseline<br>Error<br>Allowable Error | 1212 | 142<br>140<br>2<br>acceptable | 3.64<br>3.93<br>0.29<br>acceptable | 143<br>140<br>3<br>acceptable | 3.61<br>3.93<br>-0.32<br>acceptable |
| 3 | Test<br>Baseline<br>Error<br>Allowable Error | 1337 | 143<br>140<br>3<br>acceptable | 3.58<br>3.93<br>-0.35<br>acceptable | 143<br>140<br>3<br>acceptable | 3.56<br>3.93<br>-0.37<br>acceptable |
| 4 | Test<br>Baseline<br>Error | 1457 | 142<br>140<br>2 | 3.52<br>3.93<br>-0.41 | 143<br>140<br>3 | 3.50<br>3.93<br>-0.43 |
|  | Allowable Error |  | acceptable | acceptable | acceptable | acceptable |
| 5 | Test<br>Baseline<br>Error<br>Allowable Error | 1582 | 142<br>140<br>2<br>acceptable | 3.46<br>3.93<br>-0.47<br>acceptable | 144<br>140<br>4<br>acceptable | 3.46<br>3.93<br>-0.47<br>acceptable |
| 6 | Test<br>Baseline<br>Error<br>Allowable Error | 1702 | 143<br>140<br>3<br>acceptable | 3.40<br>3.93<br>**- 0.53**<br>**not acceptable** | 144<br>140<br>4<br>acceptable | 3.40<br>3.93<br>**-0.53**<br>**not acceptable** |
| 7 | Test<br>Baseline<br>Error<br>Allowable Error | 1827 | 142<br>140<br>2<br>acceptable | 3.37<br>3.93<br>**-0.56**<br>**not acceptable** | 143<br>140<br>3<br>acceptable | 3.35<br>3.93<br>**-0.58**<br>**not acceptable** |
| 8 | Test<br>Baseline<br>Error<br>Allowable Error | 1952 | 143<br>140<br>3<br>acceptable | 3.30<br>3.93<br>**-0.63**<br>**not acceptable** | 144<br>140<br>4<br>acceptable | 3.31<br>3.93<br>**-0.62**<br>**not acceptable** |

Table NO (4-4-3): Group 3 glucose concentration from 2000 to 2577 mg/dl and error in sodium, potassium.

(Allowable error Na+ 4mmol/l, K+ 0.5 mmol/l)

Table NO (4-4-3) :Group 3 glucose concentration from 2000 to 2577 mg/dl and error in sodium ,potassium. (Allowable error Na<sup>+</sup> 4mmol/l ,K<sup>+</sup> 0.5 mmol/l )
| No | Profile | Glucose conc. | Na+ direct | K+ direct | Na+ direct | K+ indirect |
| --- | --- | --- | --- | --- | --- | --- |
| 1 | Test<br>Baseline<br>Error<br>Allowable Error | 2077 | 142<br>140<br>2<br>acceptable | 3.26<br>3.93<br>**-0.67**<br>**not acceptable** | 144<br>140<br>4<br>acceptable | 3.27<br>3.93<br>**-0.66**<br>**not acceptable** |
| 2 | Test<br>Baseline<br>Error<br>Allowable Error | 2202 | 142<br>140<br>2<br>acceptable | 3.20<br>3.93<br>**-0.73**<br>**not acceptable** | 144<br>140<br>4<br>acceptable | 3.24<br>3.93<br>**-0.69**<br>**not acceptable** |
| 3 | Test<br>Baseline<br>Error<br>Allowable Error | 2327 | 142<br>140<br>2<br>acceptable | 3.15<br>3.93<br>**-0.78**<br>**not acceptable** | 144<br>140<br>4<br>acceptable | 3.20<br>3.93<br>**-0.73**<br>**not acceptable** |
| 4 | Test | 2452 | 142 | 3.11 | 144 | 3.17 |
|  | Baseline |  | 140 | 3.93 | 140 | 3.93 |
|  | Error |  | 2 | **-0.82** | 4 | **-0.76** |
|  | Allowable Error |  | acceptable | **not acceptable** | acceptable | **not acceptable** |
| 5 | Test | 2577 | 142 | 3.07 | 144 | 3.12 |
|  | Baseline |  | 140 | 3.93 | 140 | 3.93 |
|  | Error |  | 2 | **-0.86** | 4 | **-0.81** |
|  | Allowable Error |  | acceptable | **not acceptable** | acceptable | **not acceptable** |

## 5 Discussion

The measurement of ions like Na+, K+ is most commonly done by electrochemistry that is based on the measurement of an electrical signal genrated by achemical system in an electrochemical cell. Ion selective electrodes use this principle of potentiometry to measure electrolytes and are routinely used in clinical laboratories for the same.However,ISEs do not measure concentration (c);they measure only activity(a) of ions, defined as product of activity coefficient(y) and concentration (a=c x y).yis assumed to be 1 at infinite dilution and is dependent on the concentration and valence of all the ions that are present(11) In indirect ISE methods The sample is diluted with high ionic activity buffers in ratios of 1:20 – 1:34 as per the analytical systems involved [11, 12]. This ensures that the activity coefficient virtually remains constant for different samples. Thus indirect ISE yields a good estimate of concentration yielding results comparable to flame photometry. Most of the modern day autoanalyzers use this method; however they are calibrated using standard solutions with normal concentration of solids (proteins and lipids) approximately 7 % of the total plasma volume. Hence they are susceptible to ‘pseudohyponatremia’, a condition most evident in samples with hyperlipemia and hyperproteinemia [13]. On the other hand, direct ISE methods, where the electrode is directly exposed to undiluted sample are commonly available at the point of care for rapid determination of electrolytes in whole blood. It ensures measurement of the physiology-cally active fraction of the ion of interest and thus not affected by concentration of solids in the plasma (proteins and lipids). However, in biological samples like blood, plasma or serum the activity coefficient, would vary from sample to sample and should never reach beyond 0.7 under normal circumstances. Besides, variations due to changes in ionic strength due to other ions present has to be also accounted for [12]. Estimation of Na+ has been reported to be affected by several factors: the type of electrode used, the amount and type of heparin used, the pH and bicarbonate level in blood to name a few [14]. Apart from the above, some studies have suggested possible interference in Na+ estimation due to high glucose levels in the sample [5, 14]. In the present study we evaluated the effects of increasing glucose concentrations on the measurement of Na+, K+ by direct as well as indirect ISE methods.

This study is differente from Asila Al-Musheifri and Graham R D Jones(2010). When no effect in both sodium and potassium by direct and indirect methods.

This study is diffrante from Serap Cuhadar, Mehmet Koseoğlu, Yasemin Cinpolat, Guler Buğdaycı, Murat Usta, Tuna Semerci (2016). When no effect in both sodium and potassium by direct and indirect methods.

This study is agree in part of effect with Bela Goyal1 • Sudip Kumar Datta1 • Altaf A. Mir1· Saidaiah Ikkurthi1 • Rajendra Prasad1 • Arnab Pal1 (2015) When estimate sodium by indirect ion selective electrode but in correlation differ when strong negative correlation but in this study middium correlation, and diffrante in case of potassium.

## 6 Conclucion

Increasing blood gluose interferance in electrlytes mainly potasssium by both direct and indirect ion selective electrode, when increasing glucose potassium decreae significantly strong negative correlation.

No different significant between direct and in direct ion selective electrods in potassium when used cobas c311 and easy lyte.

Significant interferance in sodium by indirect ion selective electrode when used cobas c311. while no significant interferance in sodium direct ion selective electrode when used easy lyte.

Glucose concentration interfer the electrolyte specially potassium in concentration more than 1700 mg/dl by direct and indirect methods.

### 6-1 Limitation and Recommendation

All investigation done in one lab by two different devise easy lyte for direct ion selective electrode and cobas c311 for indirect ion selective electrode.

Urine for sodium and potassium not included in study because many labs not tested for rezone for not requested or to avoid contamination.

Serum and urine chloride not including in study because it is more expensive and not requested as rotine wark.

To avoid bias two or more labs in studies are recommended.

## Glucose concentration an and reading of sodium and potassium by direct and indirect ion selective electrode

**Glucose concentration and reading of sodium and potassium by direct and indirect ion selective electrode**
| **glucose conc** | **Na (direct)** | **K (direct)** | **Na (indirect)** | **K (indirect)** |
| --- | --- | --- | --- | --- |
| 110 | 140 | 3.93 | 140 | 3.93 |
| 230 | 143 | 3.98 | 144 | 4 |
| 347 | 142 | 3.94 | 142 | 3.89 |
| 467 | 142 | 3.9 | 142 | 3.85 |
| 592 | 142 | 3.88 | 142 | 3.81 |
| 717 | 142 | 3.81 | 143 | 3.77 |
| 842 | 142 | 3.76 | 143 | 3.7 |
| 967 | 141 | 3.72 | 143 | 3.67 |
| 1087 | 143 | 3.68 | 143 | 3.64 |
| 1212 | 142 | 3.64 | 143 | 3.61 |
| 1337 | 143 | 3.58 | 143 | 3.56 |
| 1457 | 142 | 3.52 | 143 | 3.5 |
| 1582 | 142 | 3.46 | 144 | 3.46 |
| 1702 | 143 | 3.4 | 144 | 3.4 |
| 1827 | 142 | 3.37 | 143 | 3.35 |
| 1952 | 143 | 3.3 | 144 | 3.31 |
| 2077 | 142 | 3.26 | 144 | 3.27 |
| 2202 | 142 | 3.2 | 144 | 3.24 |
| 2327 | 142 | 3.15 | 144 | 3.2 |
| 2452 | 142 | 3.11 | 144 | 3.17 |
| 2577 | 142 | 3.07 | 144 | 3.12 |

## Glucose interferent substance

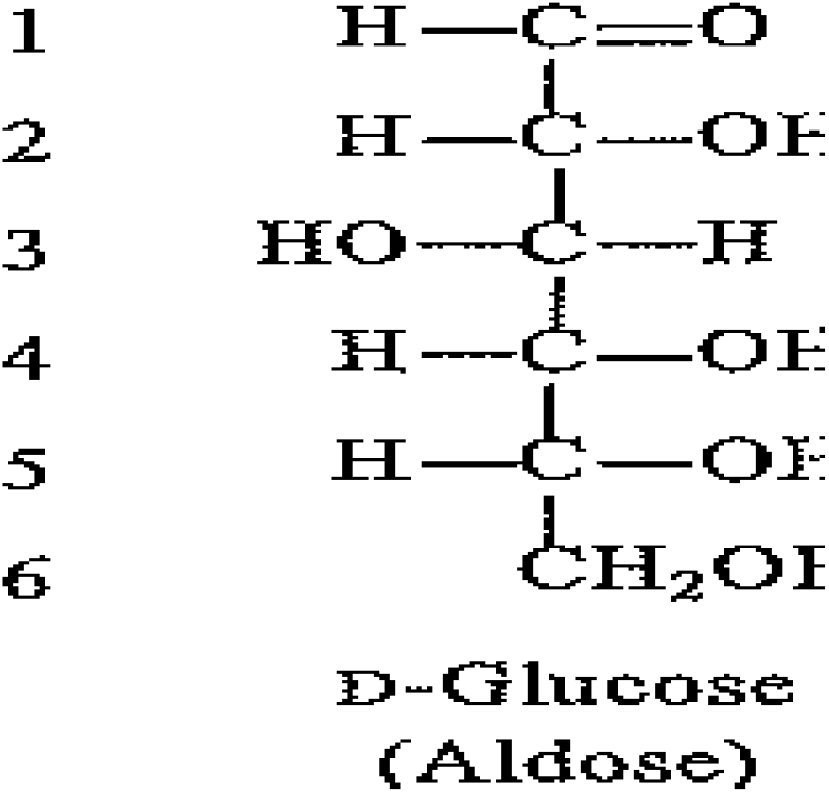

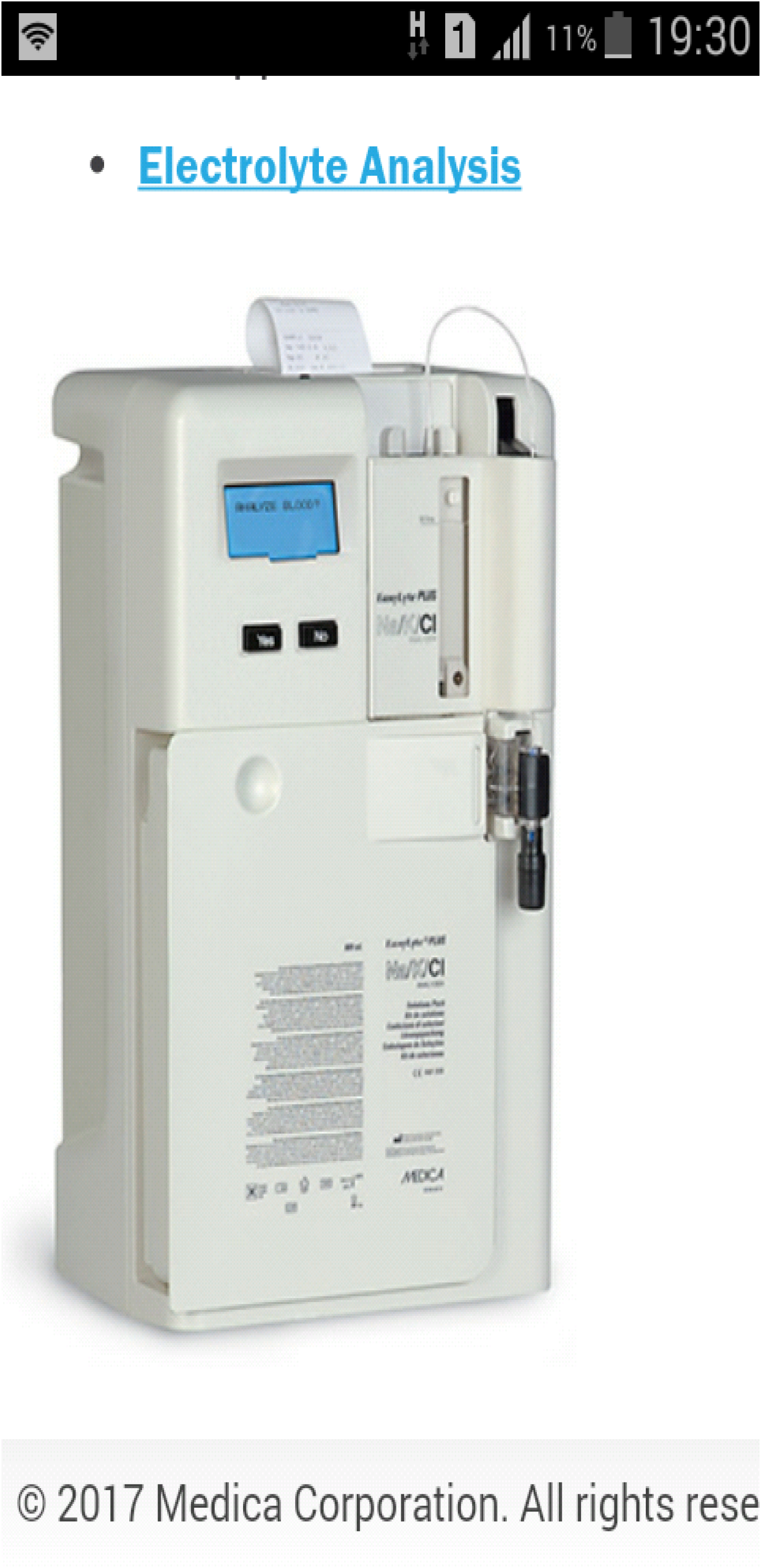

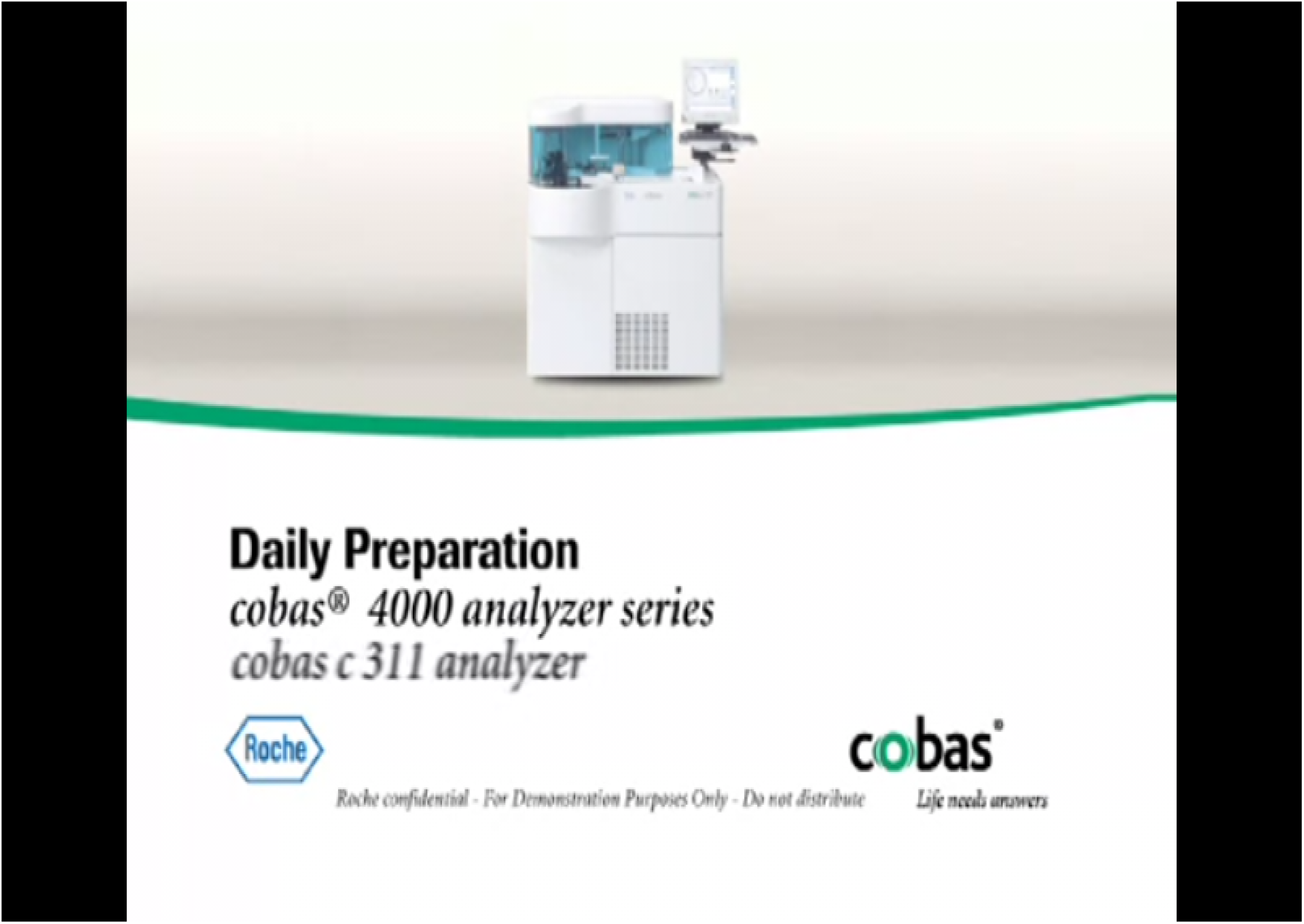

